# Multi-modal artificial dura for simultaneous large-scale optical access and large-scale electrophysiology in non-human primate cortex

**DOI:** 10.1101/2021.02.03.429596

**Authors:** Devon J. Griggs, Karam Khateeb, Jasmine Zhou, Teng Liu, Ruikang Wang, Azadeh Yazdan-Shahmorad

## Abstract

**Objective:** Non-human primates (NHPs) are critical for development of translational neural technologies because of their neurological and neuroanatomical similarities to humans. Large-scale neural interfaces in NHPs with multiple modalities for stimulation and data collection poise us to unveil network-scale dynamics of both healthy and unhealthy neural systems. We aim to develop a large-scale multi-modal interface for NHPs for the purpose of studying large-scale neural phenomena including neural disease, damage, and recovery.

**Approach:** We present a multi-modal artificial dura (MMAD) composed of flexible conductive traces printed into transparent medical grade polymer. Our MMAD provides simultaneous neurophysiological recordings and optical access to large areas of the cortex (~3 cm^2^) and is designed to mitigate photo-induced electrical artifacts. The MMAD is the centerpiece of the interfaces we have designed to support electrocorticographic recording and stimulation, cortical imaging, and optogenetic experiments, all at the large-scales afforded by the brains of NHPs. We performed electrical and optical experiments bench-side and *in vivo* with macaques to validate the utility of our MMAD.

**Main results:** Using our MMAD we present large-scale electrocorticography from sensorimotor cortex of three macaques. Furthermore, we validated surface electrical stimulation in one of our animals. Our bench-side testing showed up to 90% reduction of photo-induced artifacts with our MMAD. The transparency of our MMAD was confirmed both via bench-side testing (87% transmittance) and via *in vivo* imaging of blood flow from the underlying microvasculature using optical coherence tomography angiography.

**Significance:** Our results indicate that our MMAD supports large-scale electrocorticography, large-scale cortical imaging, and, by extension, large-scale optical stimulation. The MMAD prepares the way for both acute and long-term chronic experiments with complimentary data collection and stimulation modalities. When paired with the complex behaviors and cognitive abilities of NHPs, these assets prepare us to study large-scale neural phenomena including neural disease, damage, and recovery.

## 1. Introduction

Non-human primates (NHPs) are a critical translational model for neuroscience (Camus *et al*., 2015; Harding, 2017; Mitchell *et al*., 2018). NHP brains are evolutionarily similar to humans and they can be trained to perform complex cognitive and behavioral tasks. Traditionally, neurophysiological research techniques studying neural phenomena were limited to small brain areas and singular modalities due to technological constraints. Such constraints in turn limited the degree of insight which could be gained. However, in recent years researchers have recognized that neural computations often incorporate multiple brain regions in a wide array of phenomena including post-traumatic stress disorder (Lanius *et al*., 2015), aging (Li *et al*., 2015), memory (Huijgen and Samson, 2015; Jeong *et al*., 2015), schizophrenia (Dong *et al*., 2018), creativity (Shi *et al*., 2018), and others. Studies of large-scale brain activity often employ functional magnetic resonance imaging (fMRI), a modality with low spatial and temporal resolution. Further, fMRI is unable to perform neural stimulation – a critical tool for translational neurotherapeutic research. To increase spatial and temporal resolution and to provide stimulation capabilities at large-scales, modalities beyond fMRI must be considered in translational NHP research.

To study the cortex of NHPs at these large-scales, two key methodologies have been developed over the years: imaging (Shtoyerman *et al*., 2000; Chen *et al*., 2002, 2005; Slovin *et al*., 2002; Lu *et al*., 2010; Ruiz *et al*., 2013; Li *et al*., 2017; Ju *et al*., 2018) and electrocorticography (Bosman *et al*., 2012; Fukushima *et al*., 2015; Komatsu *et al*., 2017; Chao *et al*., 2018; Miyakawa *et al*., 2018; Chiang *et al*., 2020a, 2020b; Kaiju *et al*., 2021). To support cortical imaging, a transparent artificial dura implanted in a cranial chamber is often used to provide optical access (e.g., (Slovin *et al*., 2002)). This optical window can support months of optical access before tissue growth over the surface of the brain clouds the window. To support chronic electrocorticography, the traditional approach is to close the surgical wound after implanting an electrocorticographic (ECoG) array (e.g., (Komatsu *et al*., 2017)). Recently, a study has proposed an array integrated with an artificial dura implanted in a cranial chamber (Chiang *et al*., 2020b). This approach obscured optical access to the brain because of the high density of traces and electrodes laid across the cortex. However, we had previously demonstrated for the first time that chronic large-scale micro-ECoG and optical access to the NHP cortex could be achieved simultaneously with semi-transparent electrode arrays laid underneath an artificial dura (Yazdan-Shahmorad *et al*., 2015, 2016). Specifically, we used our interface to perform large-scale (~3 cm^2^) fluorescence imaging and micro-ECoG recording, as well as optogenetic stimulation across large cortical areas. While this setup was effective, we recognized that integrating electrodes directly into an artificial dura optimized for optical access would reduce experimental complexity.

To this end, we present a flexible, transparent artificial dura with embedded electrodes, which we call a multi-modal artificial dura (MMAD). We present both bench-side data and acute *in vivo* data from NHPs under anesthesia highlighting the multi-modal functionality of our MMAD. Our MMAD is designed to be resistant to photo-induced artifact, to be capable of simultaneous surface electrical recording and stimulation, and to provide a large-scale optical window (~3 cm^2^) to the cortex. The MMAD has long cables which allow neurophysiology systems acutely attached to the MMAD to be placed away from the optical window. This arrangement provides access for imaging hardware during experiments. The MMAD equips us for large-scale acute optical experiments in NHP cortex, such as optogenetic viral vector construct evaluation, with simultaneous electrophysiological recordings. Our MMAD is also designed to bring improved stability to our previously published chronic neural interfaces (Yazdan-Shahmorad *et al*., 2015, 2016; Griggs *et al*., 2019) which would set the stage for optimized large-scale chronic multi-modal experiments with NHPs in the future. We also outline the remaining steps to chronic MMAD implementation.

## 2. Methods

We present an MMAD designed for large-scale *in vivo* optical and electrophysiological experiments in NHPs.

### 2.1. Subjects

Three adult female pig-tailed macaques (*Macaca nemestrina*) were used (Monkey D, 12.8 kg, 14 years; Monkey E, 13.1 kg, 14 years; Monkey F, 13.8 kg, 14 years). All animal care and experiments were approved by the University of Washington’s Office of Animal Welfare, the Institutional Animal Care and Use Committee, and the Washington National Primate Research Center (WaNPRC).

### 2.2 Multi-modal Artificial Dura

Our MMAD is composed of an artificial dura made of a medical-grade transparent polymer with embedded electrodes providing surface ECoG recordings (Fig. 1a-b). Ripple Neuro (Ripple Neuro, Salt Lake City, UT) had introduced flexible, biocompatible, and semi-transparent ECoG electrodes, which we custom-designed for our MMADs. These arrays have interleaving opaque conductive layers (250-300 μm trace width with 800 μm pitch) and transparent insulating layers (50-60 μm thick) made of medical grade polymer (Fig. 1a-c). Each array provided a 20 mm diameter circle (314 mm^2^) of optical access and 32 platinum electrodes (500 μm diameter, 2mm pitch) which can be used for both electrical recording and stimulation (Fig. 1d). An extra wide polymer skirt surrounded the grid of electrodes to aid in modifying the MMAD for experiment-specific needs (Fig. 1a). The MMAD is ultra-flexible because the conductive material is composed of platinum particles dispersed in a matrix within the polymer. To protect the conductive traces and electrodes from photo-induced artifacts, an additional set of non-recording traces and electrodes were printed directly above the traces and electrodes used for electrophysiology. We refer to this additional layer as the light-blocking layer (Fig. 1d). The light-blocking layer traces used for this iteration of MMADs were printed nominally 4x wider than the underlying traces and slightly larger than the underlying electrodes. These light-blocking layer traces permit optical access to 79% of the electrode recording area and about 76% of the full optical window. The traces extend out from the array through flexible cables and terminate in contacts which serve as an electrical junction between the array and clamp-connectors (Ripple Neuro, Salt Lake City, UT), which contain PCBs (Fig 1e). These clamp-connectors are only necessary during electrophysiological experiments which protects their internal PCBs from corrosion and enables easier storage of the MMADs for chronic *in vivo* experiments. Each clamp connector has a pin for electrical grounding.

**Figure 1.**
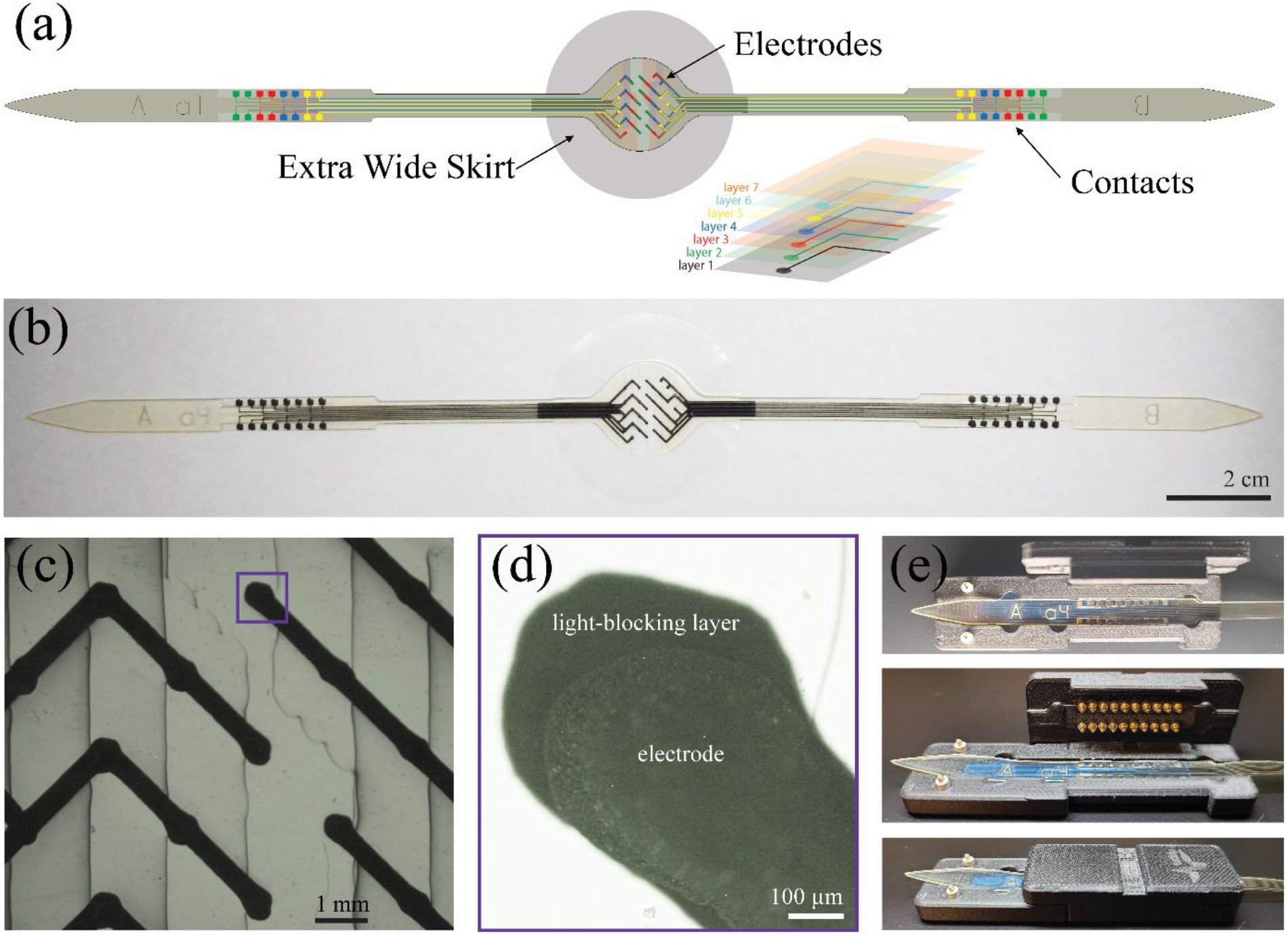
Multi-modal artificial dura (MMAD). **(a)** MMAD schematic. The MMAD is semitransparent and contains individually wired, opaque electrodes in the center with 250-300 μm traces embedded within 5 layers of a transparent medical grade copolymer. The traces extend down cables on opposite ends of the centered electrode array. The light-blocking layer (layer 1) is furthest from the cortex and mitigates photo-induced artifacts. The layered design causes the cortex to contact several layers. **(b)** MMAD. **(c)** Microscopic image of a portion of the electrode array. Edges of the polymer layers are evident. The box indicates the electrode shown in (d). **(d)** Microscopic image of one electrode (foreground) and its light-blocking layer (background). Due to the plane of focus for this image, the texture of the electrode can be observed near its edges. **(e)** MMAD cable positioned in a clamp-connector. MMAD contacts are aligned with pads on the printed circuit board.

### 2.3 Array Impedance

After production of the arrays and before implantation Ripple Neuro measured the impedance of all electrodes of four MMADs (N=128 electrodes, i.e., 4 arrays x 32 electrodes per array; measured dry with Fluke 179 True RMS Multimeter). Additionally, they measured the electrochemical impedance spectroscopy of all electrodes at 10.0 Hz, 100.0 Hz, and 1.0 kHz (electrode sites and leads submerged in normal saline solution). We performed additional impedance testing at 1 kHz with a Grapevine Nomad neurophysiology system at different timepoints post-production (Ripple Neuro, Salt Lake City, UT).

### 2.4 Optical Transmittance

We used white light to measure the optical transmittance of both the traditional transparent silicone-based artificial dura and the MMAD at five different wavelengths throughout the visible and infrared spectra (Thorlabs PM100D Power Meter and S121C Sensor). A mask with a 1 mm pinhole was used to increase the spatial specificity of the light source. We placed the beam in the center of both samples and, in the case of the MMAD, the beam of light was positioned on a transparent region (between the traces and electrodes).

### 2.5 Photo-induced Artifacts

To assess the light-blocking layer’s efficacy, we optically stimulated both sides of an MMAD with a laser (Doric, Quebec, QC) via a fiberoptic cable and recorded results. The electrodes of the MMAD as well as the tip of the fiberoptic cable were submerged in normal saline, and the fiberoptic was approximately perpendicular to the surface of the MMAD. We focused the laser, pulsing at 1 Hz and at 1 ms per pulse, on one electrode while simultaneously recording from the array. Then we flipped the MMAD to illuminate the electrode from the light-blocking layer side, which replicates the orientation during *in vivo* experiments, and repeated the illumination and recording process. We performed this entire process for two neighboring electrodes and for both green (520 nm) and red light (638 nm; data not shown). We also performed this entire process focusing the laser near, but not at, the same electrodes. The optical power of the laser was approximately 51 mW for green light and 104 mW for red light as measured with a photodiode (respectively measured at 535 nm and 635 nm; Thorlabs PM100D Power Meter and S121C Sensor).

### 2.6 Bench-side Electrical Stimulation

To verify our MMAD’s capability of performing simultaneous stimulation and recording, we applied bench-side electrical stimulation at a single electrode with all electrode sites submerged in normal saline. We used a stimulation train with 5 Hz biphasic pulses and a 200 ms pulse width. The stimulation amplitude was 10uA for both phases. During each 1-min stimulation session we recorded from all 32 sites on the MMAD.

### 2.7 Connector holder

We designed a connector holder to secure the MMAD on the cortical surface and to hold the clamp-connectors during acute, anesthetized experimentation (Fig. 2a). An intra-cranial rim on the bottom of the connector holder fit inside a craniotomy (see section 2.8). This fit aided in keeping the connector holder stationary as holes were drilled for the skull screws necessary to affix the connector holder to the skull. The connector holder was 3D printed with plastic filament (acrylonitrile butadiene styrene (ABS) with quick support release (QSR) used as supports) which provided flexibility as the fastening of the skull screws warped the connector holder slightly against the curved skull surface. Slots on either side of the cranial window allow the flexible cables of the MMAD to exit the craniotomy and extend to the clamp-connectors (Fig. 2). The clamp-connectors were secured to the arms of the connector holder with rubber bands which hooked onto the crossbars of the arms. The angle of the arms was chosen to provide optimal clearance between the clamp-connectors and the sterile field, while still providing room for imaging equipment. The connector holder also included screw posts to aid in affixing or aligning additional experimental equipment to the connector holder, such as optical stimulation devices we have presented previously (Yazdan-Shahmorad *et al*., 2015, 2016; Griggs *et al*., 2019), although these posts were not used in the experiments presented here.

**Figure 2.**
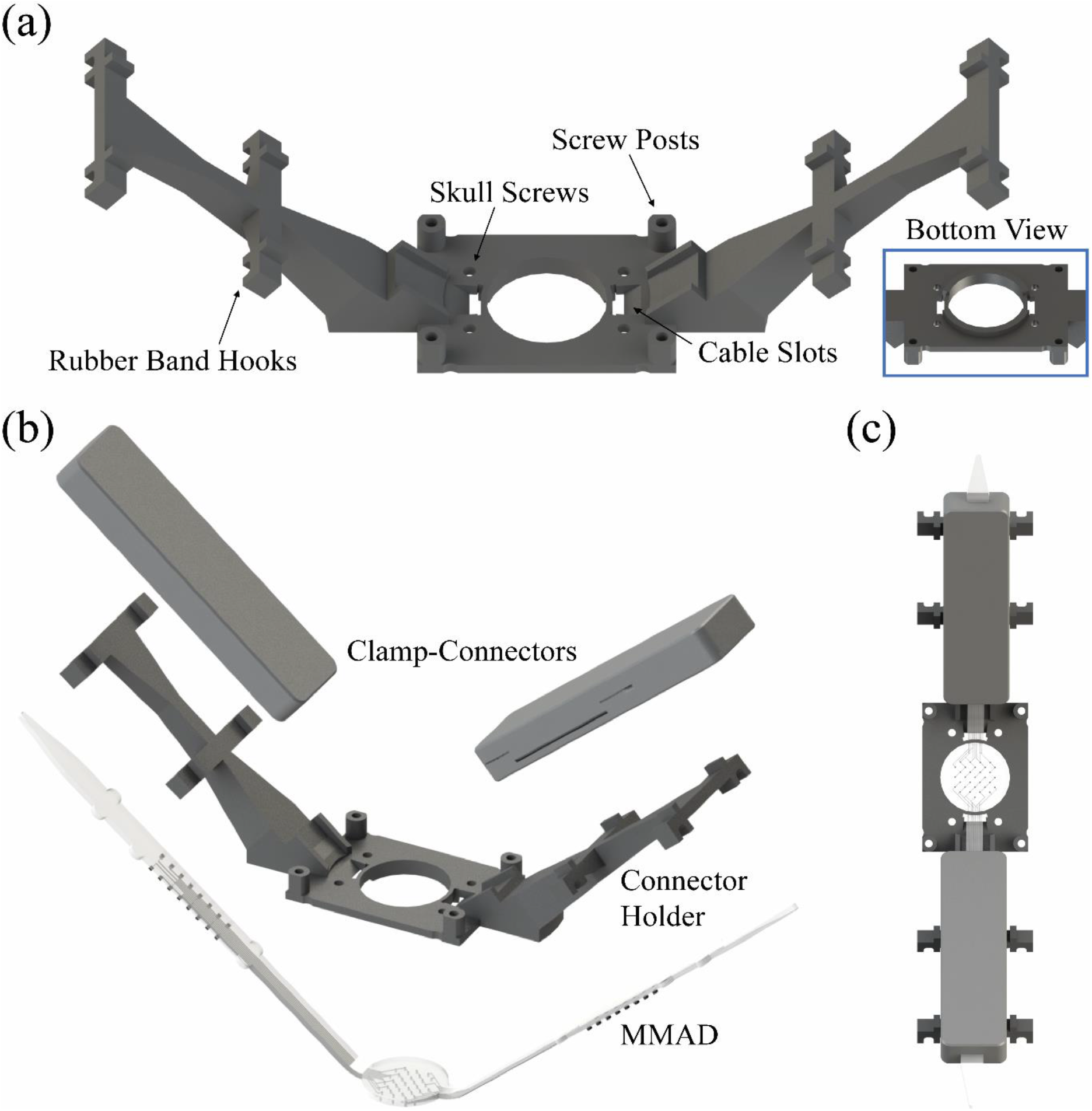
Acute interface. **(a)** Acute interface connector holder. The connector holder secures the MMAD electrode array against the brain and holds the clamp-connectors (b) away from the subject with rubber bands which attach to the rubber band hooks. The cables of the MMAD are passed through the cable slots to interface with the clamp-connectors (b, c). The connector holder is secured to the skull with skull screws inserted through holes near the cranial window. Additional equipment may be affixed or aligned to the connector holder with the screw posts. Inset: Bottom view of the connector holder revealing the intra-cranial rim which aids in centering the MMAD and connector holder over the craniotomy. **(b, c)** Acute interface stack-up. The MMAD array cables thread through the slots and are secured in the clamp-connectors which rest on either arm of the connector holder.

### 2.8 Surgical Preparation

We collected data from all subjects during terminal experiments. The subject’s temperature, heart rate, and respiration were monitored throughout the procedure. To begin surgical preparations, the subject was anesthetized with isoflurane and mounted into a stereotactic frame. An incision was made down the midline of the head. The skin and musculature were peeled from the skull laterally with elevators to reveal both sides of the skull. Then we used a trephine to create bilateral 25 mm diameter craniotomies targeting the primary motor and primary somatosensory cortices. We performed a durotomy by raising the dura off of the cortex with a needle and cutting out the dura with ophthalmic scissors. We cut four notches in the extra wide skirt, two adjacent to each MMAD cable, then we placed the MMAD on the cortex. The notches permitted us to insert the extra wide skirt under the skull and native dura while still allowing the cables to extend out of the cable slots in the connector holder. Connector holders were affixed to the skull bilaterally with screws to hold the MMADs in place and support the clamp-connectors. Each connector holder includes four skull screw holes, however only the two medial screw holes for each connector holder were necessary in practice. The arms of the connector holders extended roughly parallel to the midline so that the connector holders would not interfere with each other. We implanted an additional skull screw near the midline and away from the craniotomies to serve as an electrical ground – we connected conductive wires from the grounding pins of each clamp connector to the screw. The subjects were transitioned to urethane anesthesia (1 g/kg dose) prior to electrophysiology recordings and supplemental doses were administered as required to maintain the level of anesthesia.

### 2.9 Electrophysiology

We recorded ECoG data at 30 kHz sampling frequency bilaterally from all three subjects for 30 minutes and performed electrical stimulation in the left hemisphere of Monkey F. All electrophysiology data collection and electrical stimulation were performed with a Grapevine Nomad and Nano front ends (Ripple Neuro, Salt Lake City, UT).

### 2.10 *In Vivo* Electrical Stimulation

We implanted two MMADs bilaterally as described in section 2.8. We performed electrical stimulation delivered through a single electrode on the MMAD implanted on the left hemisphere of Monkey F. Stimulation took place in blocks after 30 minutes of baseline recording. Each of six stimulation blocks lasted 10 minutes, with 2 minutes of baseline recordings in between the blocks to track changes in electrophysiology. The stimulation trains had a 5 Hz burst frequency with five 1 kHz biphasic pulses within each burst. The stimulation amplitude was set to 65 μA for both phases.

### 2.11 Optical Coherence Tomography Angiography

To validate the transparency and imaging capabilities of our MMAD *in vivo*, we performed optical coherence tomography angiography (OCTA) through the MMAD placed on the cortex of the right hemisphere of Monkey D. Imaging and image processing were performed with previously described methods (An *et al*., 2010; Deegan *et al*., 2018). We used an articulating arm to position the OCTA probe over the optical window. The low profile of the portion of the connector holder near the craniotomy provided access for the OCTA probe to be positioned near the MMAD. Electrophysiological data were not collected during this proof-of-concept imaging session. Of note, current OCTA techniques require post-processing which prevents real-time visualization of large-scale images.

## 3. Results

As described above, we validated our MMAD both with bench-side tests and acute *in vivo* experiments.

### 3.1. Bench-side Data

#### 3.1.1. Array Impedance

The direct current resistance of the electrodes had a mean of 342.7 Ω and a standard deviation of 21.4 Ω, and all electrodes had an isolation resistance higher than 10 MΩ. The electrochemical impedance spectroscopy of all electrodes at 10.0 Hz, 100.0 Hz, and 1.0 kHz respectively had means of 5.56 MΩ, 1.59 MΩ, and 268.8 kΩ and respective standard deviations of 1.59 MΩ, 0.31 MΩ, and 46.3 kΩ (Fig. 3a).

**Figure 3.**
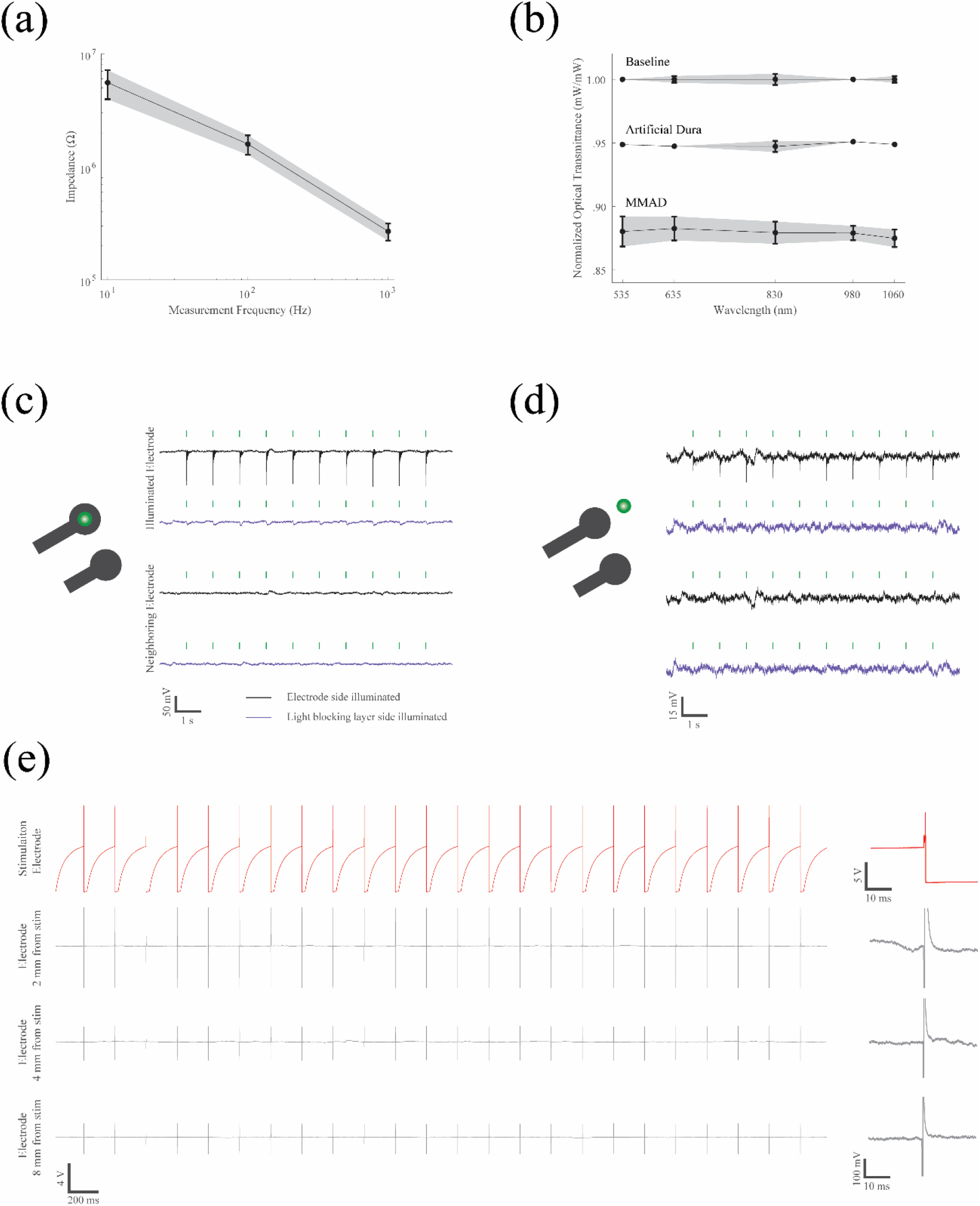
Bench-side results. **(a)** Initial mean impedance and standard deviation of four MMADs with 32 electrodes each (total electrode count N=128) measured at 10 Hz, 100 Hz, and 1 kHz. **(b)** Mean and standard deviation optical transmittance of a classic silicone-based artificial dura and our MMAD. A white light source was projected through a mask with a 1 mm pin hole which, in the case of the MMAD, was aligned to project between electrode traces. Results were recorded at five wavelengths within the visible and infrared spectra. **(c)** Photo-induced artifacts of an electrode illuminated by green light, and a neighboring electrode recorded simultaneously. Black traces indicate when the electrode was illuminated directly (i.e., the MMAD was upside down) and purple traces indicate when the electrode’s light-blocking layer was illuminated. Both the tip of the fiberoptic cable and the MMAD were submerged in saline. **(d)** Photo-induced artifacts as in (c) where the MMAD was illuminated near, but not on, the electrode. **(e)** Electrical stimulation waveforms. Left panel: Stimulation channel (red) and other recording channels (grey) at varying distances away from the stimulation channel. Right panel: Example stimulation waveforms for each channel.

We used the same two MMADs for all three acute experiments: the first two acute experiments were conducted 5 to 6 months post array production while the third experiment was conducted 15 months post production. Both arrays demonstrated bench-side impedance values within the expected range through the first experiment. However, during and after the third *in vivo* experiment, we found 10 failed electrodes with abnormally high impedance and electrically disconnected recording traces on each array. Further bench-side testing of two unused arrays with the same production date also showed 12 and 8 bad electrodes respectively. Upon further investigation, we found that these failed electrodes were largely printed on the same conductive layer across all 4 arrays.

#### 3.1.2. Optical Transmittance

When normalized, the optical transmittance values of the artificial dura and MMAD resulted in approximately 95% and 88%, respectively, across all five wavelengths measured (Fig. 3b). The variance in the MMAD data is greater than that of the artificial dura. We believe the variance in optical properties due to the multi-layering approach of MMAD fabrication accounts for the variation in transmittance.

#### 3.1.3. Photo-induced Artifacts

For both electrodes and both light colors we found that the light-blocking layer reduced photo-induced responses by 87–90% (Fig. 3c-d, responses measured peak-to-peak). Since the photo-induced response was limited to the stimulated electrode, we can infer that the induced artifact was caused by the illumination as opposed to electrical noise. The illumination process was repeated with the laser directed near, but not at, the electrode. In this case we found that the light-blocking layer practically eliminated photo-induced artifact. We did not observe any difference between red and green illumination.

#### 3.1.4. Stimulation Waveforms

During the bench-side electrical stimulation, we observed rapid signal recovery within 10 ms at all recording channels except the stimulation channel (Fig. 3e). When comparing raw stimulation waveforms at 2 mm, 4 mm, and 8 mm away from the stimulation electrode, we also found that the amplitude of stimulation artifact and signal recovery time both decreased with the distance between our recording and stimulation sites. This fast signal settling suggests that our MMAD is able to simultaneously stimulate and record neural activity across a large area surrounding the stimulation site.

### 3.2 Acute *In Vivo* Data

We validated surface electrical recording and stimulation, as well as cortical imaging through our MMAD *in vivo*.

#### 3.2.1 Optical Coherence Tomography Angiography

We demonstrated the ability to perform OCTA imaging through the semi-transparent MMAD. Our acute setup permitted us to position imaging hardware close to the MMAD surface. Importantly, the transparency of the MMAD allowed for imaging of blood flow in the underlying microvascular structures down to 600 μm in depth, which is the maximum achieved with OCTA under the current experimental setting (Fig. 4). As expected, the electrodes and traces occluded the OCTA field of view. However, the extent of disruption was minimal and did not prevent imaging in the semi-transparent regions of the MMAD. The data collected through the MMAD were qualitatively similar to data collected through a transparent silicone-based artificial dura (data not shown).

**Figure 4.**
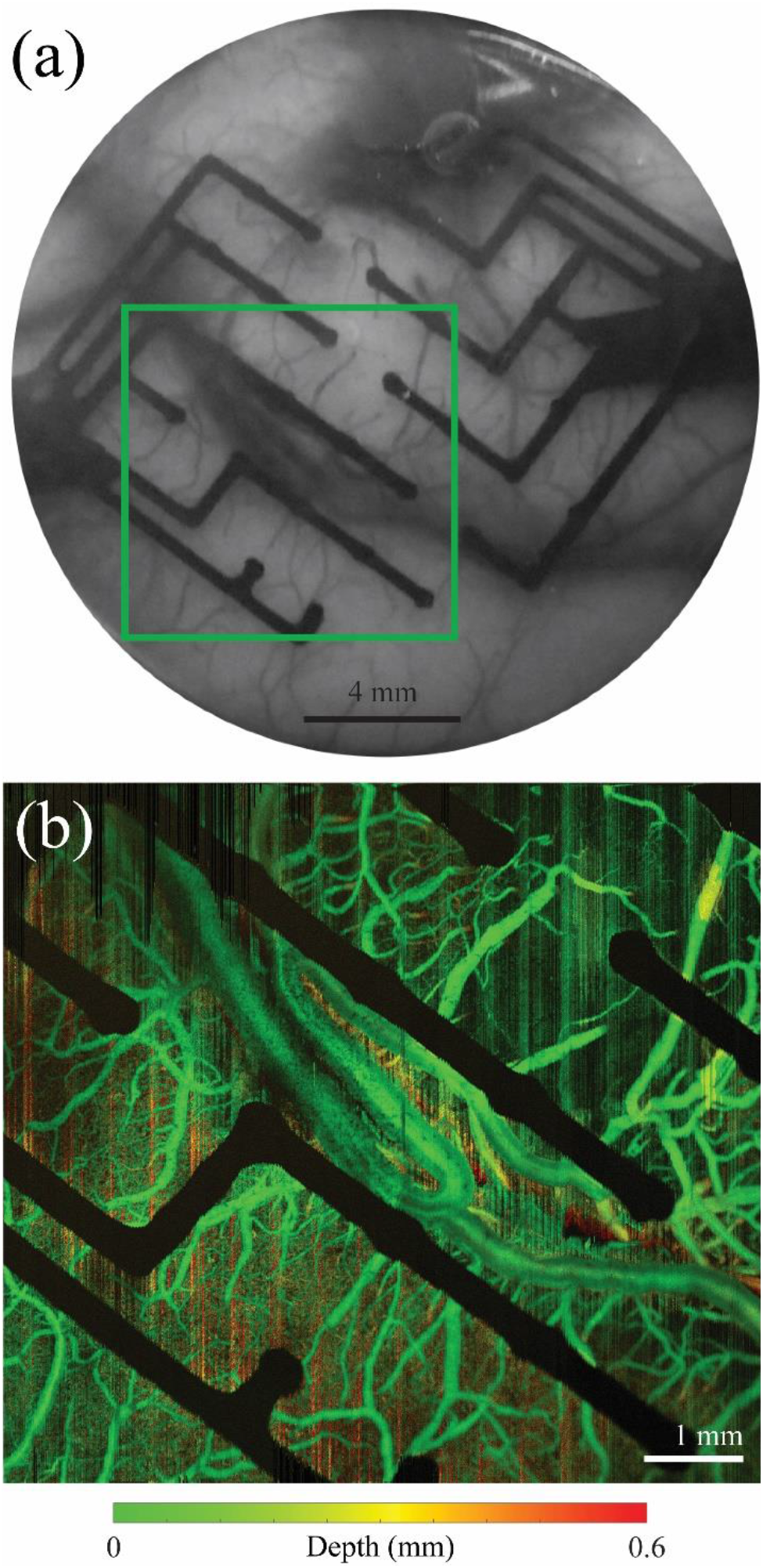
*In vivo* imaging. **(a)** Greyscale picture of the MMAD on the sensorimotor cortex of an NHP. Box indicates the region imaged in (b). **(b)** Optical coherence tomography angiography of the cortex as imaged through the array.

#### 3.2.2 ECoG Recordings

We successfully collected bilateral ECoG data from all three subjects under urethane anesthesia (Fig. S1). The power spectral densities of the electrodes calculated over a 30-minute period approximate a 1/*f*^α^ curve, while peaks are evident at 60 Hz and its harmonics, as expected (Fig. 5a). Example heatmaps from bilateral recordings of Monkey D in different frequency bands reveal a heterogenous distribution of power and groupings of electrodes (Fig. 5b). During recording, we observed artifacts when the animal was touched. In addition, across the two MMADs implanted in each subject, we saw two bad channels for Monkey D, four bad channels for Monkey E, and 20 bad channels for Monkey F with abnormally low signal power, which were excluded from analysis. Further testing suggested that the bad channels we observed from Monkey F recordings matched the failed electrodes we found in bench-side impedance measurements (see section 3.1.1).

**Figure 5.**
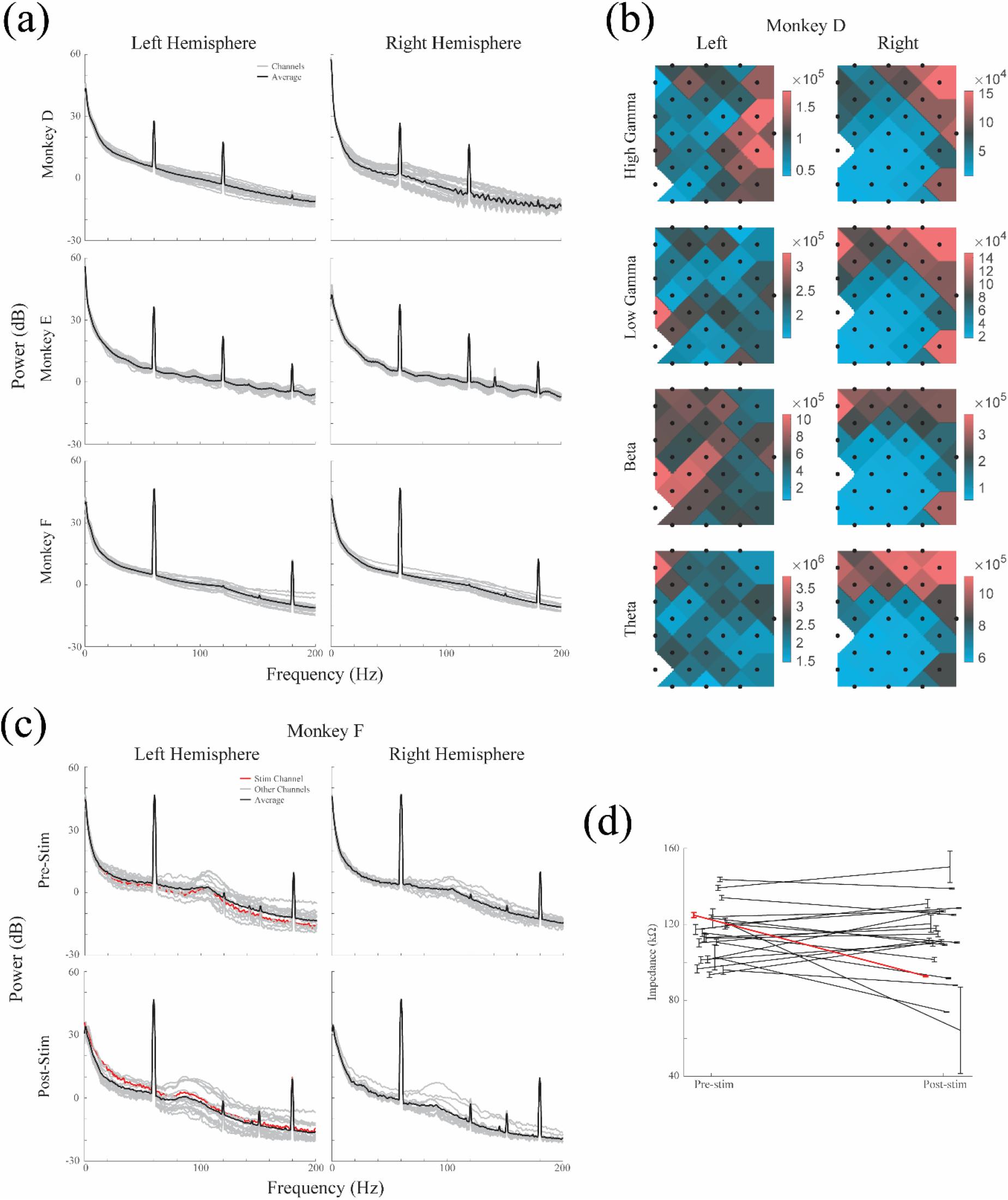
*In vivo* electrocorticography and stimulation. **(a)** Power spectral densities of left and right hemispheres of three NHPs, recorded with MMADs for 30 minutes at 30 kHz sampling frequency. Individual electrodes are traced in grey and the mean is traced in black. Power lines cause noise spikes at 60 Hz and its harmonics. High impedance electrodes have been omitted. **(b)** Electrode array power heatmaps of left and right hemispheres of Monkey D at four frequency bins: theta (4–7 Hz), beta (12–29 Hz), low gamma (30–59 Hz), and high gamma (60–150 Hz). In each array one electrode had high impedance and was omitted. **(c)** Power spectral densities of left and right hemispheres of Monkey F recorded for two minutes before and after 20 minutes of stimulation. Individual electrodes are traced in grey, the stimulating electrode is traced in red, and the mean is traced in black. High impedance electrodes have been omitted. **(d)** Standard error of the mean impedance measurements of electrodes before and after stimulation. The electrode used for stimulation is shown in red.

#### 3.2.3 ECoG Stimulation

We successfully performed electrical stimulation on the left hemisphere of Monkey F while recording bilateral electrophysiology. We calculated the power spectral densities both before and after stimulation and observed a 1/*f*^α^ curve in both cases which is characteristic of neural data (Fig. 5c, left column). The power spectral densities compared between the stimulated and non-stimulated hemispheres were qualitatively similar as well (Fig. 5c). Due to reduced recording time (2 minutes) the traces from before and after stimulation (Fig. 5c) are expected to be less smooth than the baseline recording traces (Fig. 5a). Specifically, we present the power spectral densities after 30000 stimulation pulses (20 minutes) to give us a reliable perspective of stimulation impact on electrodes.

We measured the impedance of the electrodes before and after stimulation with our electrophysiology system’s built-in 1 kHz impedance measurement tool. To prevent bad channels from skewing the data, we omitted any impedance greater than 850 kΩ. We found that impedances did change for some electrodes (Fig. 5d; mean: −2.2 kΩ, −1.9%; standard deviation: 18.7 kΩ, 16.3%). However, when the distribution of the mean impedances of the electrodes were compared between before and after stimulation with a paired t-test, no significance was found (p=0.60). Similarly, the stimulated electrode did not exhibit impedance changes statistically different from the other electrodes (p=0.055). The success of electrical stimulation and recording from MMADs shows efficient electrical connection between our MMAD and our recording system and validates our design which eliminates the need for permanent attachment of PCBs to the MMAD. This shows the efficacy of our design and paves the path for chronic implementation of our MMADs.

## 4. Discussion

Artificial duras have been used for decades to provide large-scale optical access to the brains of NHPs (Arieli *et al*., 2002; Slovin *et al*., 2002; Chernov and Roe, 2014; Nassi *et al*., 2015; Seidemann *et al*., 2016; Yazdan-Shahmorad *et al*., 2016; Zaraza *et al*., 2020). Over the years there have been advances in large-scale optical imaging and optical stimulation modalities, including the discovery (Boyden *et al*., 2005) and widespread proliferation of optogenetics (Yizhar *et al*., 2011). However, there has been little change in artificial dura designs for NHPs. Here, we present a multi-modal artificial dura, being an artificial dura with embedded electrodes. The MMAD is designed with multiple layers of traces and electrodes vertically aligned in a transparent polymer to increase the transparent surface area, making it well fit for large-scale optical modalities. We have previously demonstrated micro-ECoG arrays which supported large-scale optogenetics in NHPs (Yazdan-Shahmorad *et al*., 2016), but the arrays were fragile and not suitable to be molded into an artificial dura. In contrast, our MMAD can be utilized as both an electrode array and an artificial dura because our conductive traces are flexible. Additionally, our previous micro-ECoG arrays were attached to PCBs which were prone to corrosion. In our present design, the absence of permanently attached PCBs makes the MMAD setup corrosion resistant. The wide spacing between the electrode traces would provide easy integration with penetrating electrical or optical probes similar to others (Ruiz *et al*., 2013). Moreover, combining an artificial dura and an electrode array by printing flexible traces and electrodes directly into the artificial dura curtails experimental complexity by reducing the number of components required for multi-modal experiments.

Traditional artificial duras have been used in NHPs for a variety of optical imaging and stimulation modalities including intrinsic optical imaging (Arieli *et al*., 2002; Ruiz *et al*., 2013; Chernov and Roe, 2014; Zaraza *et al*., 2020), voltage sensitive dye imaging (Arieli *et al*., 2002; Slovin *et al*., 2002), calcium imaging (Seidemann *et al*., 2016), fluorescence imaging (Nassi *et al*., 2015; Yazdan-Shahmorad *et al*., 2016), OCTA (Khateeb *et al*., 2019b), infrared neural stimulation (Chernov and Roe, 2014), and optogenetics (Ruiz *et al*., 2013; Nassi *et al*., 2015; Yazdan-Shahmorad *et al*., 2016). With our MMAD we aim to accommodate optical modalities such as these in tandem with ECoG recording. After characterizing the MMAD with optical and electrical bench-side tests, we validated the utility of our MMAD via an acute implementation by 1-collecting bilateral ECoG data from the cortex of three NHPs, 2-performing electrical stimulation in one of the subjects, and 3-performing OCTA imaging through the MMAD in one of the subjects. Analysis of the acute data presented in this paper show that the MMAD effectively performs both ECoG recording and stimulation while maintaining optical access for OCTA. Electrophysiological data were not collected during our proof-of-concept imaging session. However, we do not expect to have any issues collecting electrophysiology data while performing OCTA or other large-scale optical modalities such as optogenetics and calcium imaging in NHPs. In particular, our OCTA data show we can image the blood flow through the array and our previous optogenetic data (Yazdan-Shahmorad *et al*., 2015, 2016) show that as long as the light sources are not directly placed on top of an opaque electrode or trace, light penetration would be sufficient to evoke reliable neural responses.

A novel aspect of our design is the technology developed by Ripple Neuro (Salt Lake City, UT) which provides the capability to attach and detach the PCBs from the MMADs through clamp-connectors. This enables us to store the MMADs chronically in cranial chambers, reduce the risk for corrosion of the electronics, and reduce the complexity of chamber design. The electrical stimulation and recording capabilities demonstrated here validate the electrical connection between the electrodes and our recording system through the clamp-connectors.

Concerns of generating photo-induced artifacts naturally arise when optical and electrophysiological modalities are combined. The most obvious approach to avoiding photo-induced artifacts is directing the light away from metal traces. For optogenetic applications, we can also use opsin kinetics to distinguish between light-evoked neural activity and photo-induced artifact similar to our previous work (Ledochowitsch *et al*., 2015). In that work, we stimulated at frequencies higher than our opsin off-kinetics. If light-evoked signal was detected from electrodes near the stimulated area, we concluded that signal was non-biological. This was only happening if the tip of our fiberoptic was directly located on an electrode and moving the fiber away slightly was solving the problem. We can use this technique to identify photo-induced artifacts in our MMAD as well.

In this paper, we also demonstrate that the addition of a light-blocking layer over electrodes reduces photo-induced artifacts substantially when optical stimulation is directed at the electrode, and practically eliminates artifact when the optical stimulation is directed near, but not at, the electrode. Like the recording traces, the light blocking layer traces are printed with platinum particles because of their strong adhesion properties. As mentioned above, when the fiberoptic was placed directly above the electrode, we observed reduced photo-induced artifact through the light-blocking layer. It is worth noting that the artifact waveform in this case was shaped distinctly different from the artifact waveform produced without the light-blocking layer. In particular, the former has a slow rise and fall, while the later has a sharp onset and quickly falls off. This slow response suggests some capacitive effects between the light-blocking layer and the electrode. To address this issue in future MMAD iterations, we plan to electrically ground the light-blocking layer or print the light-blocking layer with a highly opaque non-conductive material. We also plan to vary the width of the light-blocking layer traces to optimize the balance between artifact protection and optical access to the brain. Our present MMAD design eliminates photo-induced artifact when the light is not directed at the electrode. For instances when a light source is close to the electrode site we can use the artifact detection process described above to mitigate the artifact. In sum, our data suggest that the MMAD is suitable for simultaneous optical and electrophysiological modalities. We plan to demonstrate such simultaneous data collection in future work.

Given that all MMAD electrodes tested immediately before the first *in vivo* experiment showed normal impedance values, the two bad channels we saw from Monkey D recording were likely caused by poor contact between the MMAD tails and PCB clamp connectors, not intrinsic electrode failure. However, from the *in vivo* experiment in Monkey F and the subsequent bench-side impedance testing, we observed that some of the electrodes from the same print layer on both the used and unused MMAD arrays went bad between 6 and 15 months post production. We are working with Ripple Neuro to understand the major reasons underlying these failures and improve our next batch of arrays being produced. However, even with these limitations, our bench-side and *in vivo* data still suggest that we can reliably use the MMAD in animal experiments for at least 6 months after production.

Additional limitations of our MMAD are the number, size, and pitch of electrodes. These properties arise from a combination of current printing constraints and trade-offs between electrode coverage and optical access. Another related limitation is that cells directly beneath the electrodes are shielded from optical modalities. However, we have previously shown that optogenetic stimulation near an electrode can reliably evoke neural responses which can be recorded by opaque electrodes (Yazdan-Shahmorad *et al*., 2016). In the future, we plan to investigate alternative fabrication methods to provide greater electrode array coverage without compromising optical access and flexibility.

While other groups have developed interfaces for large-scale imaging (Shtoyerman *et al*., 2000; Chen *et al*., 2002, 2005; Slovin *et al*., 2002; Lu *et al*., 2010; Ruiz *et al*., 2013; Li *et al*., 2017; Ju *et al*., 2018), large-scale electrocorticography (Bosman *et al*., 2012; Fukushima *et al*., 2015; Komatsu *et al*., 2017; Chao *et al*., 2018; Miyakawa *et al*., 2018; Chiang *et al*., 2020a, 2020b; Kaiju *et al*., 2021), and large-scale optogenetics (Komatsu *et al*., 2017; Rajalingham *et al*., 2020) in NHPs, and still others have proposed a variety of multi-modal systems with other animal models (Kwon *et al*., 2012; Kuzum *et al*., 2014; Richner *et al*., 2014; Ji *et al*., 2017, 2018; Liu *et al*., 2018; Stamatakis *et al*., 2018; Zátonyi *et al*., 2018; Renz *et al*., 2020), to our knowledge we remain the only group to have brought all three technologies together chronically for NHPs (Yazdan-Shahmorad *et al*., 2015, 2016). This suggests that a simpler and more robust interface may be broadly advantageous. With this in mind, we designed our MMAD to be adapted for chronic use.

Our previous interface consisted of a chronically implanted chamber but required daily implantation and explantation of the micro-ECoG array, which was a time-consuming process (Yazdan-Shahmorad *et al*., 2016). Additionally, this process introduced unwanted variance in day-to-day electrode positions on the cortex, and daily perturbation of the cortex appeared to accelerate tissue growth on the surface of the brain, obstructing optical access in 4-5 weeks. In order to restore optical access we had to wait for the tissue to mature and separate from the cortex so that the tissue could be resected for continued optical experiments (Yazdan-Shahmorad *et al*., 2016). When we tried chronically implanting micro-ECoG arrays beneath the artificial dura, we observed accelerated tissue growth over the top of the array, which obstructed optical access within a few weeks, thus also forcing us to wait to perform a resection (Yazdan-Shahmorad *et al*., 2015). Due to these issues with both acute and chronic implementation of the micro-ECoG arrays in conjunction with artificial duras, we designed our MMAD to prevent tissue growth over the top of the electrode array during chronic implantation by printing the electrode array into the artificial dura itself. Based on our past experience with traditional artificial duras (Yazdan-Shahmorad *et al*., 2016) and similar experiences of others (Arieli *et al*., 2002; Chen *et al*., 2005; Ruiz *et al*., 2013), we expect that an artificial dura with integrated electrodes, as opposed to a separate electrode array, will reduce tissue growth on top of the brain while maintaining large-scale optical and electrophysiological access to the brain. Furthermore, bringing optical access and electrodes together into a single device would reduce experimental complexity. We previously published designs for a cranial chamber to chronically stabilize the MMAD on the cortex (Griggs *et al*., 2019). We expect this stabilization to mitigate variance in day-to-day electrode positioning and reduce perturbations of the cortex which, as previously stated, appeared to accelerate tissue growth on the surface of the brain (Yazdan-Shahmorad *et al*., 2016). The cranial chamber is also designed to house the flexible, PCB-free, corrosion-resistant MMAD cables in a coiled configuration between experiments as we have shown bench-side (Griggs *et al*., 2019) which will eliminate our previous corrosion issues with permanently mounted PCBs (Yazdan-Shahmorad *et al*., 2015).

We outlined a roadmap to chronic implantation of our MMAD in Fig. S2. We have completed bench-side and acute in vivo experiments in this work, and we have previously published designs for a chronically implantable chamber (Griggs *et al*., 2019). In the future, we plan to implant the chamber, perform a viral vector transfusion with methods similar to those we have previously published (Yazdan-Shahmorad *et al*., 2016; Khateeb *et al*., 2019a), and implant the MMAD, molded into its chronic form (Griggs *et al*., 2019), in a single surgery. We expect that viral expression will be sufficiently pronounced roughly 2 months after the surgery, at which point we will be prepared to perform chronic in vivo experiments with electrophysiology, electrical stimulation, imaging, and optogenetic modulation.

Our MMAD in combination with our previously presented chronic interfaces (Yazdan-Shahmorad *et al*., 2015, 2016; Griggs *et al*., 2019) equip us for a variety of research pursuits. Moving forward, we plan to capitalize on our previous optogenetic (Ledochowitsch *et al*., 2015; Yazdan-Shahmorad *et al*., 2015, 2016, 2018b, 2018a, 2018c; Khateeb *et al*., 2019a; Ojemann *et al*., 2020; Tremblay *et al*., 2020) and imaging (Khateeb *et al*., 2019b; Macknik *et al*., 2019) experience as we pioneer the experimental spaces unveiled by the merging of large-scale ECoG and large-scale optical access to the NHP cortex. The synergistic power of these technologies poise us to build on our past work investigating neural plasticity (Yazdan-Shahmorad *et al*., 2018a; Bloch *et al*., 2019) as we study large-scale, chronic neural phenomena including neural disease, damage, and recovery. To specifically mention one example research pursuit, we plan to use the MMAD to study the effects of stimulation during recovery from our photothrombotic stroke model (Khateeb *et al*., 2019b).

## 5. Conclusion

In this paper, we are integrating electrodes into a traditional artificial dura, expanding its functionality. Our presented MMAD for NHPs supports both large scale optical methods and large-scale electrical recording and stimulation. Specifically, we presented acute *in vivo* data demonstrating the utility of our MMAD for OCTA imaging and electrophysiological recording and stimulation of the large brains of macaques. In addition, our MMADs are compatible for chronic, large-scale electrophysiological experiments complimented by optical techniques, such as imaging and optogenetics.

## 6. Acknowledgments

We would like to thank Toni Han and WaNPRC staff for helping prepare for and run the surgeries; WaNPRC staff for animal care; William Ojemann, Julien Bloch, and Maryam Bahadori-Nejad for helping with interface design; and Minh Nhan Le for helping with the OCTA collection. We would also like to thank Alex Thiessen, Jessi Mischel, and Jose Ortega from Ripple Neuro for their help with the design and manufacturing of the MMADs.

None of the authors have any conflicts of interest to disclose.

This project was supported by the Eunice Kennedy Shriver National Institute of Child Health & Human Development of the National Institutes of Health (K12HD073945, AY), the University of Washington Royalty Research Fund (AY), the Washington National Primate Research Center (WaNPRC, P51 OD010425), the Center for Neurotechnology (CNT, a National Science Foundation Engineering Research Center, EEC-1028725, DG), the National Science Foundation Graduate Research Fellowship (KK), and the Weill Neurohub (JZ).

AY conceptualized the experiments, AY and RK provided supervision and resources; AY, RK, and KK acquired funding; DG, JZ, KK, and AY designed experiments; DG, JZ, KK, TL, and AY performed data collection; JZ, KK, TL, and AY performed data analysis; DG wrote the original draft of the manuscript; DG, JZ, KK, TL, RK, and AY revised and edited the manuscript.

---

**Supplementary Figure 1.**
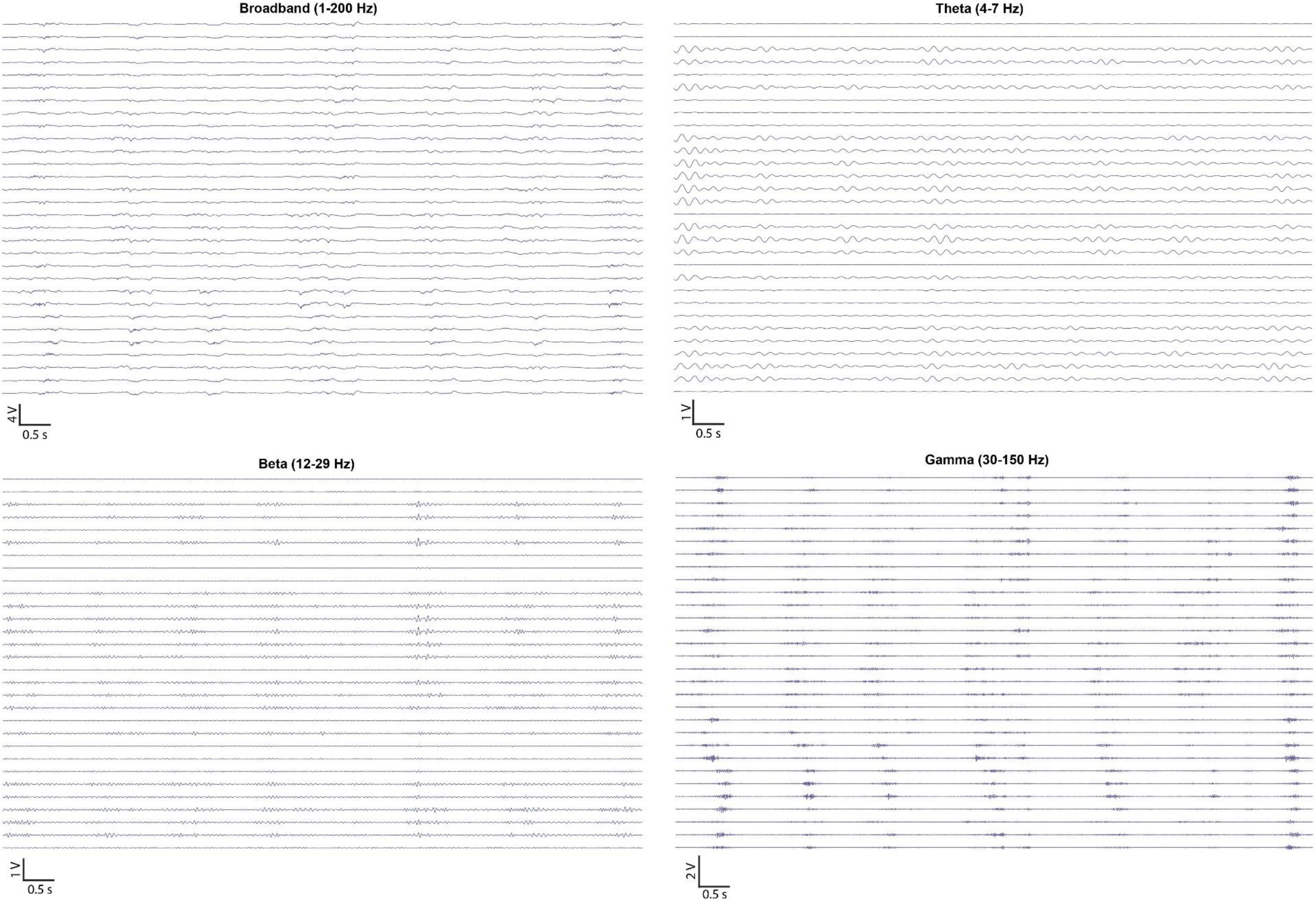
Example *in vivo* electrocorticography time series filtered to different frequency bands.

**Supplementary Figure 2.**
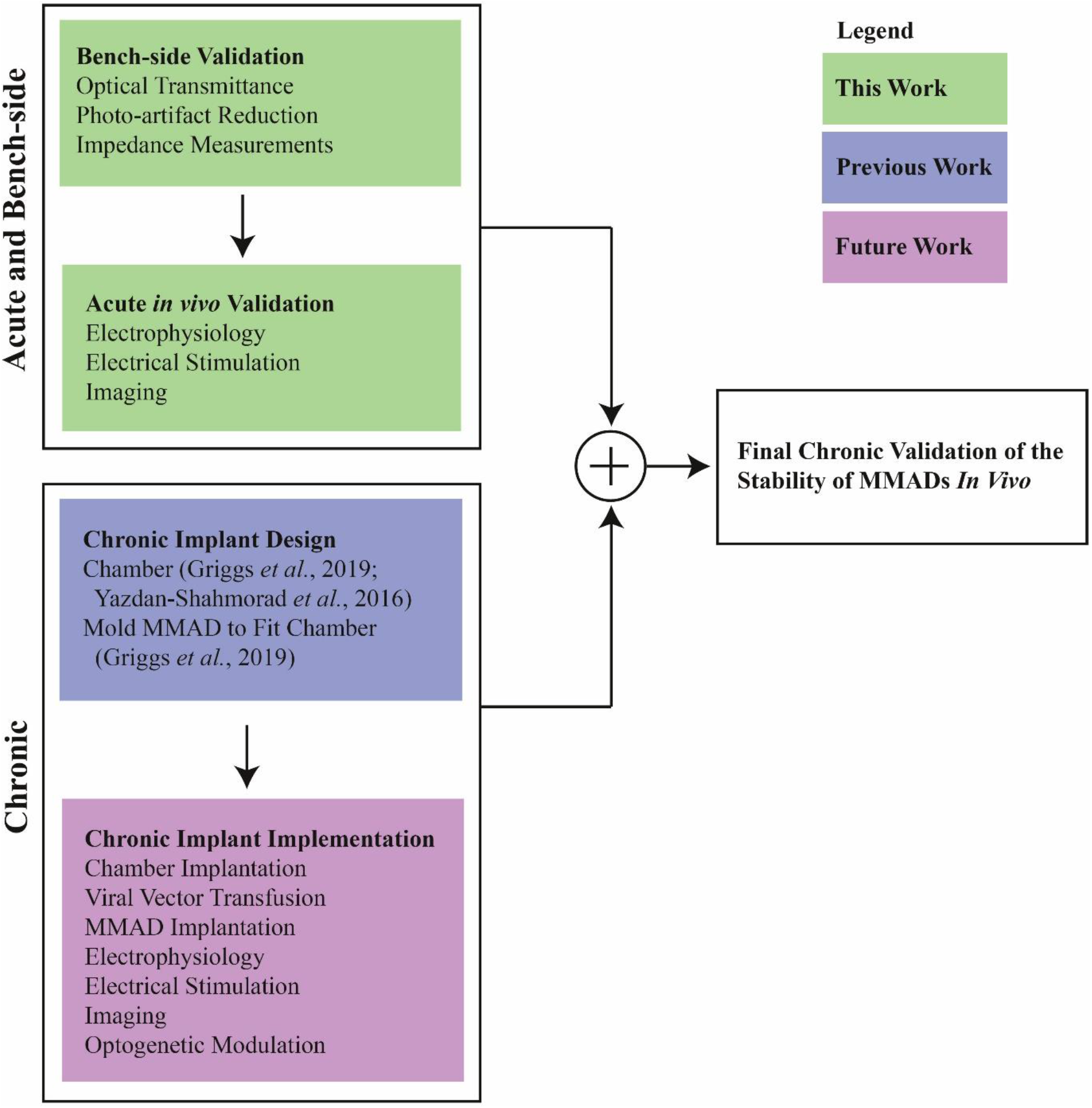
Roadmap to chronic validation of MMAD.

